# Vacuum dressing of collective water response and biological hydration

**DOI:** 10.64898/2026.11.25.768838

**Authors:** N. Haramein, W. Brown

**Affiliations:** ISFS SA, Martigny, Switzerland

## Abstract

Electromagnetic boundaries can modify the collective vibrational response of water. We examine how this response could influence hydrogen-bond organization and biological hydration. A Green-tensor framework relates boundary-dependent field fluctuations to the reaction field acting on water polarization and distinguishes vacuum dressing from external excitation. In an idealized infrared resonator, tuning the cavity to the O–H resonance at fixed pump input increased the calculated collective vibrational amplitude 2.69-fold for the chosen parameters, below resolved polariton splitting. An anharmonic coupling links the mean-square O–H displacement to the force on a hydrogen-bond coordinate. In a separate experiment, we acquired duplicate Fourier-transform near-infrared (FT-NIR) transmission spectra of each of two seawater-derived saline samples before and after a 15-min powered electromagnetic treatment. After multiplicative scatter correction, both samples showed changes in the same direction at 11 of 12 water matrix coordinates. Mean corrected absorbance increased by 1.07 and 1.65 milli-absorbance units near 1490 and 1512 nm, respectively, consistent with greater relative contributions from more extensively hydrogen-bonded or strongly bound water. These bands provide spectral targets for testing how electromagnetic perturbations influence water at biological interfaces.

## 1 Introduction

Water is both a molecular solvent and a responsive component of biological organization [1]. Its intermolecular hydrogen bonds couple local structure to vibrational spectra, dielectric response, and the hydration of macromolecules [2]. The possibility of controlling that response through the electromagnetic environment therefore raises a specific biological question: could modified collective water polarization influence the hydration conditions that shape protein behavior?

Existing work provides a basis for this question. Solvent-slaving models relate protein conformational motions to bulk-solvent and hydration-shell fluctuations, with dipolar interactions proposed as a mechanism linking solvent dynamics to charged protein groups [3, 4]. Molecular simulations predict strongly polarized hydration shells and non-Gaussian electrostatic fluctuations at protein–water interfaces [5]. Broadband dielectric spectroscopy, combined with simulations, resolves distinct hydration-water responses around bovine serum albumin and indicates a dynamical influence extending beyond the first hydration layer [6]. These studies support coupling between collective water response and the environment experienced by a protein.

A direct perturbation study reported that approximately 0.1 THz irradiation accelerated hydration changes in initially unequilibrated lysozyme solutions, with dielectric and NMR observations interpreted as a shift toward hydrophobic hydration distinguishable from heating controls [7]. The direction and extent of coupling nevertheless depend on the system: peptide simulations found that collective correlations account for much of the hydration-water dielectric retardation, while peptide polarization influenced water more strongly than the reverse [8]; static-field neutron-scattering measurements found no appreciable enhancement of hydration-water diffusion or change in the measured internal protein dynamics under the tested low-temperature conditions [9]. These precedents motivate testing an electromagnetic route to altered hydration, while leaving the causal link to a specific protein function to be established. They concern orientational and intermolecular dynamics; extending the connection to the O–H vibrational response considered here requires an explicit molecular coupling.

Engineered resonators establish that the O–H manifold of liquid water can hybridize with electromagnetic modes. Vibrational strong coupling has been demonstrated in Fabry–Pérot cavities and in structures supporting surface-lattice resonances [10, 11]. These systems provide experimentally accessible examples of collective light–matter response. Electrically tunable spectroscopy of nanoconfined crystalline water supplies a complementary precedent for linking local fields, hydrogen bonding, and O–H observables within a calibrated molecular model [12]. Together, these results motivate investigation of electromagnetic dressing under conditions extending beyond conventional optical cavities.

Our central hypothesis is that a suitable electromagnetic environment can modify collective water response and thereby influence hydrogen-bond organization, with possible consequences for biological hydration. Material boundaries establish the field propagation and reaction properties. External energy can excite the resulting response or maintain a material configuration in which those properties differ. A large static field, a driven oscillation, and a strongly hybridized mode describe different aspects of this mechanism; their relationship can be evaluated within one response framework.

The biological motivation is particularly relevant to energized interfaces, where membrane bias, composition, and geometry coexist with hydration water. We consider these interfaces as candidate environments for electromagnetic dressing rather than assume that they reproduce an engineered infrared cavity. Aquaphotomics provides a complementary experimental approach: water absorbance patterns report how aqueous systems respond to perturbation [13, 14]. This makes the near-infrared spectrum a useful target for a molecular model connecting vibration to hydrogen bonding.

We develop that connection in three steps. First, a partitioned Green tensor defines the field spectrum and polarization reaction kernel. Second, an idealized powered resonator demonstrates the amplitude and phase of a collective O–H response, followed by a minimal molecular coupling to a slower structural coordinate. Third, preliminary paired FT-NIR measurements identify a spectral pattern consistent with altered hydrogen-bond environments. The biological discussion then places these results within the broader hypothesis of collective water organization influencing hydration and function.

## 2 Green tensor and collective water response

### 2.1 Electromagnetic environment and reaction field

We retain the active O–H response explicitly and assign the remaining linear electromagnetic response to a background relative permittivity *ϵ*_*env*_. This partition prevents the same vibrational oscillator from being counted both inside the environment and again as the water subsystem. With the Fourier convention *e*^−*iωt*^, the retarded Green tensor of a nonmagnetic background obeys [15–17]

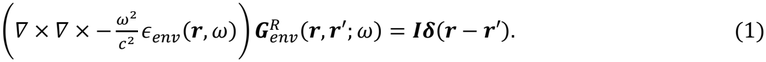

Here, ***r*** and ***r*′** denote the observation and source positions, respectively; *ω* is the angular frequency, and *c* is the speed of light in vacuum. The dimensionless, generally complex quantity *ϵ*_env_(***r***, *ω*) is the background relative permittivity, excluding the active O–H response treated explicitly. Its frequency dependence and imaginary part describe dispersion and absorption, respectively. The operator ∇×∇× takes the curl twice, with spatial derivatives acting on the observation coordinate ***r***. The identity tensor *I* and three-dimensional Dirac delta *δ*(***r*** − ***r′***) specify a point source in each Cartesian direction. The superscript ^*R*^ denotes the retarded, or causal, Green tensor: the field response cannot precede its source. The solution satisfies the electromagnetic interface conditions and outgoing-wave conditions at infinity.

The Green tensor describes the field produced at one position by a source at another. It contains the effects of boundaries, polarization, dispersion, and absorption. Its imaginary part determines the electric-field spectral density sampled by a dipole. Equation (1) fixes the electromagnetic response kernel; the field state enters through its fluctuation statistics. For a reciprocal equilibrium background, define a one-sided symmetrized spectrum whose positive-frequency integral gives the equal-time covariance:

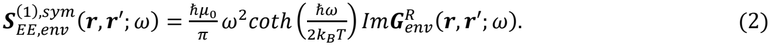

Projection along an O–H transition dipole selects the relevant polarization channel. This is the connection to the local density of electromagnetic states: a boundary can redistribute the frequencies and polarizations to which water couples. The zero-temperature limit retains zero-point fluctuations. Equation (2) specifies equilibrium fluctuations of the background; a driven nonequilibrium state requires its own source statistics. Section 2.3 makes the vacuum limit and its contribution to the dark polarization covariance explicit.

The complex Green tensor also determines the field returned by the environment to the active water polarization. For polarization density ***p***, the regularized reaction operator is

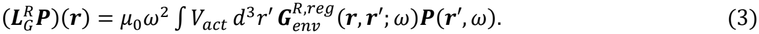

The coincident field requires a finite-volume or local-cavity prescription applied consistently to the reference and modified environments. If the molecular response has already been calibrated in bulk water, only the additional boundary reaction should be added. The imaginary part of the tensor describes fluctuation and dissipative contributions, whereas its full complex value supplies both dispersive and dissipative feedback.

### 2.2 Driven polarization and fluctuations

Let 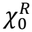 map local electric field to polarization density, with SI units F m^−1^. The incident field contains an imposed mean drive and zero-mean fluctuations. Closing the local field through Equation (3) gives

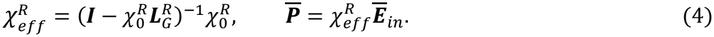

The product 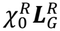is dimensionless. In a scalar projection this becomes the familiar susceptibility divided by a feedback denominator. Where the bare susceptibility is invertible, the full inverse response is 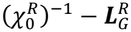. Its complex-frequency zeros identify candidate response poles, with stable poles below the real axis and a nonzero residue required in the measured channel. This formulation retains molecular resonances even when the reaction increment vanishes.

For a stable linearization, the polarization covariance follows from the same response operator:

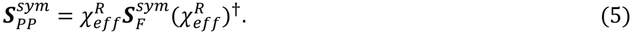

Here ***S***_*F*_ includes electromagnetic and molecular-bath fluctuations expressed as equivalent incident fields, with cross-correlations retained when required. External pumping changes the mean polarization. In a fixed linear system, an ideal coherent pump leaves the connected fluctuation covariance at its undriven value. A pump-dependent covariance or kernel therefore requires a change in the material response, noise, or nonlinear dynamics. Vacuum dressing and supplied excitation enter one electromagnetic description while retaining their distinct roles.

Collective response means that a shared field channel couples contributions from multiple molecules. Phase-referenced motion under a pump can occur below resolved strong coupling. Establishing self-sustained spatial order would additionally require nonlinear stability and correlation-length information. The present framework identifies the quantities needed for that larger investigation without making a static ordering assumption.

### 2.3 Vacuum fluctuations and the dark limit

In a dark environment with no incident drive, the O–H response still couples to electromagnetic vacuum fluctuations. Darkness alone does not remove thermal fluctuations; the ideal vacuum limit additionally takes the background field and its material reservoirs to zero temperature. To separate these contributions, write the thermal factor in Equation (2) as [16]

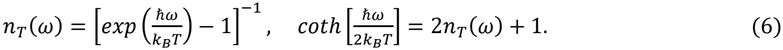

Here *n*_T_ is the thermal occupation number, *ℏ* is the reduced Planck constant, *k*_B_ is Boltzmann’s constant, and *T* is absolute temperature. The unity term is the zero-point contribution. Thus, with the one-sided symmetrized convention of Equation (2), the positive-frequency vacuum spectrum is

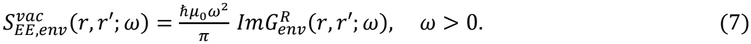

Here μ_0_ is the vacuum permeability and Im denotes the imaginary part. This spectrum remains nonzero when the thermal occupation vanishes. In an absorbing environment, it describes the electric-field correlations of the coupled background field–matter ground state. Boundaries and material response therefore reshape the vacuum correlations sampled by an O–H dipole; they need not supply incident photons. The retarded reaction kernel and dressed susceptibility in Equations (3) and (4) also remain defined without a pump.

If the electromagnetic and molecular input noises are independent and expressed in the same equivalent-field convention, the electromagnetic vacuum contribution to Equation (5) is

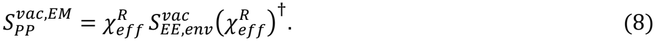

The dagger denotes the Hermitian adjoint; position integrations and any required projections are implicit as in Equation (5). This source contribution is not the full coupled ground-state covariance. Molecular reservoirs also contribute zero-point fluctuations, and all noise terms must be consistent with the dissipative response and the partition used in Section 2.1. When the input noises are correlated, their cross terms must be retained rather than assigning an independent additive vacuum contribution.

For a stable linearization about equilibrium, the undriven induced mean polarization vanishes while its connected covariance remains nonzero. Coupling can modify resonance frequencies, the decay of an initially excited vibration, and ground-state correlations. A passive stationary system in its coupled ground state does not undergo continuous excitation by absorbing vacuum photons. Upward and downward transition rates require ordered quantum correlations; the symmetrized spectrum alone is not a measure of excitation energy available to the oscillator [16].

To test a structural consequence, compare the full dark polarization covariance in the modified and reference environments using the same molecular normalization and local-field regularization. A molecularly resolved response or a justified local-coordinate projection is needed to obtain the change in local O–H displacement variance. Through the anharmonic coupling in Equation (17), that variance change can alter the mean force on the slower hydrogen-bond coordinate. A single bright-mode amplitude does not determine every local bond variance, and the electromagnetic source term in Equation (8) alone does not establish a structural change.

## 3 Idealized powered infrared resonator

### 3.1 Coupled field and vibration

The idealized resonator is motivated by the proton-cavity model of Haramein, Alirol and Guermonprez [18], which links confined vacuum-field correlations to screening boundaries and poloidal–toroidal circulation. We propose an infrared application of this principle: engineered geometry and electromagnetic response reshape the zero-point spectrum sampled by the water O–H stretch, with the aim of enhancing resonant field–dipole coupling. This spectral selection is expressed here through the Green tensor and reaction kernel, with the illustrative parameters specified below.

Consider a powered source coupled to a lossy infrared resonance overlapping water. The resonator geometry is summarized by its frequency, linewidth, mode profile, and molecular overlap. We use a single electromagnetic mode and one collective bright O–H mode, meaning the weighted combination of molecular oscillators that couples to that field. This reduction gives a nonproprietary example with specified response parameters; it is not a reconstruction of the experimental treatment apparatus that was used in the experimental investigation reported in subsequent sections of this study.

Let *a* and *B* denote dimensionless mean amplitudes of the field and bright vibrational modes. In the rotating-wave approximation, their equations are

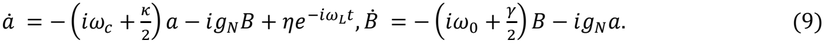

The angular frequencies *ω*_*c*_, *ω*_0_, and *ω*_*L*_ are the resonator, vibration, and pump frequencies. The uncoupled spectral full widths, in angular-frequency units, are *k* and *γ*, and *η* has units s^−1^. Quantum Langevin noise terms with zero mean accompany these equations when calculating fluctuations. The effective vibrational linewidth represents the broad liquid-water response rather than a separately measured microscopic coherence rate.

Writing 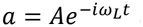 and 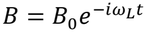, and defining *D*_*c*_ = *k*/2 − *i(ω*_*L*_ − *ω*_*c*_) and *D*_v_ = γ/2 − *i*(*ω*_*L*_ − *ω*_*0*_), gives

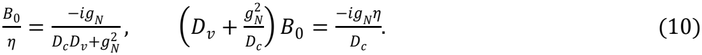

The term 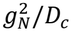is the modal reaction contribution obtained by eliminating the field. It is the reduced counterpart of the Green-tensor kernel in Equation (4). The right-hand side is the pump delivered through the resonator. This separation shows explicitly how field-mediated feedback and external excitation determine the same driven response.

The collective coupling contains the zero-point field coefficient of the mode:

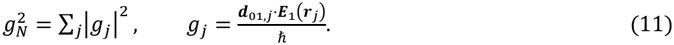

Transition dipoles, molecular orientation, and spatial overlap determine which water molecules contribute. In a nondispersive mode-volume approximation, the field coefficient is 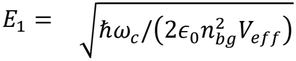 absorbing media require a consistent dispersive normalization. The pump populates the coupled system, while Equation (11) defines its vacuum coupling scale. Increasing pump amplitude alone does not increase *g*_*N*_ in this linear model.

### 3.2 Numerical response

We set the O–H center to 3400 cm^−1^, corresponding to approximately 102 THz and 2.94 µm, and choose 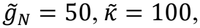 and 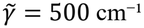. A tilde denotes division of an angular frequency or rate by 2*πc*, using c in cm s^−1^. The resulting cooperativity,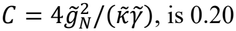. The pump is assumed to overlap the infrared mode; its existence and frequency are model inputs, independent of the FT-NIR measurements.

Figure 1 illustrates how the amplitude and phase of the collective O–H vibration depend on the driving frequency and the resonator frequency. The curves are calculated from the exact steady-state solution in Equation (10). Both panels share the same horizontal coordinate, the pump detuning from the uncoupled O–H vibrational center:

**Figure 1.**
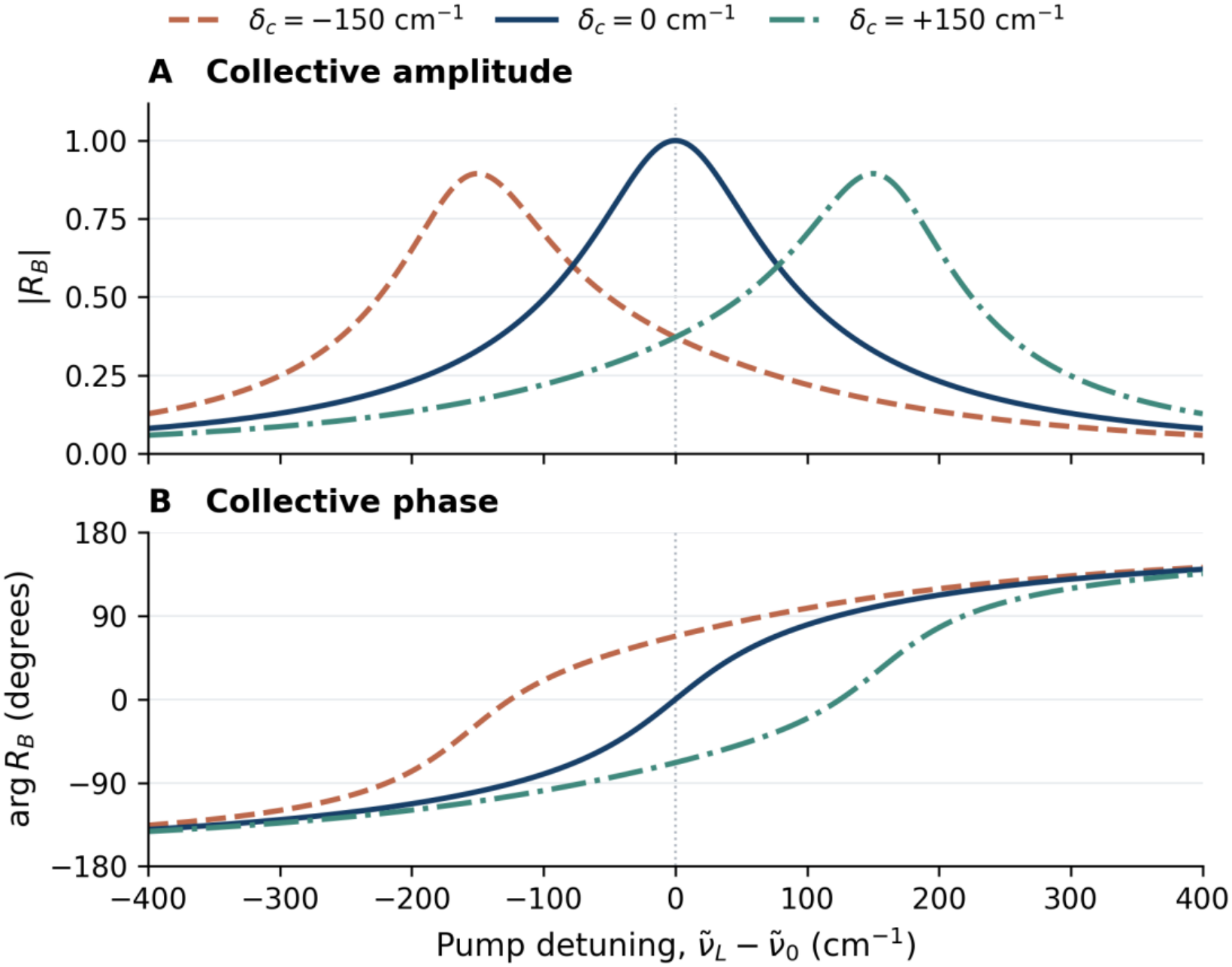
Driven collective O–H amplitude and phase in an idealized infrared environment. Exact steady solutions of Equation (10) at fixed pump amplitude, with 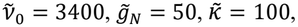and 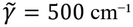. Cavity detuning is 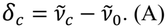. (A) Collective amplitude normalized by the response at simultaneous pump– cavity–vibration resonance. (B) Phase relative to that same reference, whose absolute phase relative to *η* is −90°. All curves use one shared normalization and unchanged coupling and losses. The vertical line marks the O–H center. The calculation demonstrates control of a driven response below resolved strong coupling; it does not fit the FT-NIR observations.

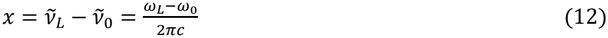

Here, 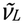 is the pump wavenumber, 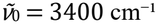, and *c* is expressed in cm s^−1^. Negative and positive detunings correspond to driving frequencies below and above the O–H center, respectively. The horizontal sweep from −400 to +400 cm^−1^ therefore spans pump wavenumbers of 3000–3800 cm^−1^. Each curve represents a fixed cavity detuning, 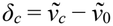 chosen as −150, 0, or +150 cm^−1^. Thus, movement along a curve changes the pump frequency, whereas comparison between curves changes the cavity frequency. The driving coefficient *η*, collective coupling, and linewidths remain fixed.

The complex amplitude *B*_0_ contains both the magnitude and phase of the collective vibrational response. To compare the three resonator settings, we normalize this amplitude by its value at simultaneous pump–cavity–vibration resonance, evaluated at the same *η*. Equation (10) then gives

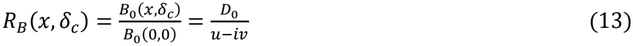

For an explicit calculation of the plotted ordinates, the real quantities *D*_0_, *u*, and *v* are defined by

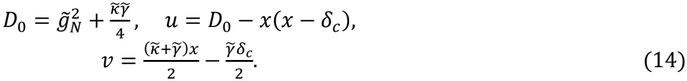

The tildes on the coupling and linewidths denote division by 2*πc*, so these quantities have the same wavenumber units as the detunings. This common normalization gives *R*_*B*_(0, 0) = 1 and preserves amplitude differences between the curves. Panel A plots the magnitude of the normalized response:

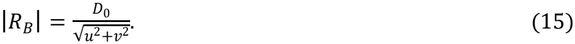

This ordinate is a dimensionless amplitude ratio. For example, a value of 0.5 corresponds to half the collective vibrational amplitude of the resonant reference. Panel B plots the phase relative to that reference, expressed in degrees:

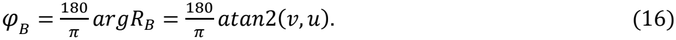

The two-argument arctangent retains the correct phase quadrant. The zero of this phase scale corresponds to the resonant reference, whose phase relative to the driving coefficient *η* is −90° under the time convention used in Equation (9).

For the selected parameters, at the O–H center (*x* = 0), the resonant cavity gives |*R*_*B*_| = 1 and *φ*_*B*_ = 0°. The cavities detuned by −150 and +150 cm^−1^ both give |*R*_*B*_| ≈ 0.3714, with relative phases of +68.2° and −68.2°, respectively. Consequently, tuning either detuned cavity to resonance while holding the pump at the O–H center increases the collective amplitude by a factor of approximately 2.69. These differences arise from excitation delivered through the resonator together with field-mediated feedback. They illustrate a tunable driven response under the chosen model parameters and do not separately quantify a change in vacuum fluctuations.

The response already changes substantially without resolved real-frequency pole splitting. At zero detuning the two poles have imaginary parts −63.4 and −236.6 cm^−1^ and the same real part. As a comparison within the same equations, increasing the chosen coupling to 300 cm^−1^ gives C = 7.2, a real pole separation of 565.7 cm^−1^, and a damping-derived linewidth of 300 cm^−1^ for each pole. This illustrates the transition toward strong hybridization as coupling competes with losses. Neither parameter set is fitted to the treatment device or interpreted as a measured hydrogen-bond energy.

Resolved strong coupling and ultrastrong ground-state dressing involve different comparisons: coupling relative to linewidths and coupling relative to bare transition frequencies, respectively. De Liberato showed that, for dissipative Lorentzian resonances, a substantial ground-state virtual-photon population can persist even when losses prevent resolved strong coupling [19]. Broad linewidths therefore do not by themselves exclude vacuum dressing, although a sufficiently large coupling relative to the transition frequency remains necessary for ultrastrong effects. This motivates examining vacuum correlations in lossy water environments. The rotating-wave calculation above illustrates driven response below resolved splitting; it does not quantify ground-state dressing or establish vacuum-induced structural changes in water. Setting the pump to zero in Equation (9) yields the undriven mean amplitudes, not the coupled ground-state covariance. A calculation of ground-state dressing must retain the terms required by its coupling regime and gauge, including counter-rotating interactions and the corresponding quadratic self-polarization or diamagnetic terms where applicable [20].

### 3.3 Coupling to hydrogen bond organization

A connection from vibration to molecular organization requires an additional structural degree of freedom. For a local O–H displacement q and a slower hydrogen-bond coordinate Q, consider the local potential expansion

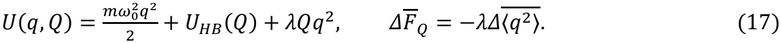

The force difference is evaluated at fixed Q, averaging over the fast vibration. Since ⟨*q*^2^⟩ = ⟨*q*⟩^2^ + ⟨(δ*q*)^2^⟩, both driven motion and fluctuations can change this mean force. In return, Q changes the local vibrational restoring coefficient, providing a molecular route for feedback between vibration and hydrogen-bond configuration. The expansion is local; higher-order terms are needed for a globally stable potential. The sign of λ, the structural restoring dynamics, and vibrational relaxation determine which configurations are favored.

The normalized bright amplitude is not itself a local bond displacement, so Equation (10) cannot fix this structural change without molecular participation factors and the remaining vibrational modes. Near-infrared overtones additionally require mechanical or electrical anharmonicity. These ingredients identify a calculable pathway toward water organization while leaving its direction to molecular modeling and experiment. Energy redistribution through pumped polaritons has, for example, been demonstrated in simulations of liquid methane [21]; its application to the hydrogen-bond network of water is a separate question.

## 4 Preliminary FT NIR observations

### 4.1 Measurement and analysis

On 4 August 2026, we measured two samples (S1 and S2) of a seawater-derived saline solution before and after powered electromagnetic treatment (Figure 2). For each sample, we introduced the saline through the inlet of a fixed flow-through cuvette, recorded two baseline transmission spectra, and withdrew the liquid through the outlet tubing. The samples were transferred to a separate apparatus for a 15-min exposure. Immediately after return, we reintroduced each sample into the same cuvette and recorded two further spectra. The cuvette remained in the spectrometer throughout. The approximately 45 min between paired acquisitions included transfer, treatment, and return. All four baseline spectra preceded the four post-treatment spectra. Mean sample temperature rose from 29.84 to 30.14 °C over this interval. The design included no matched sham treatment.

**Figure 2.**
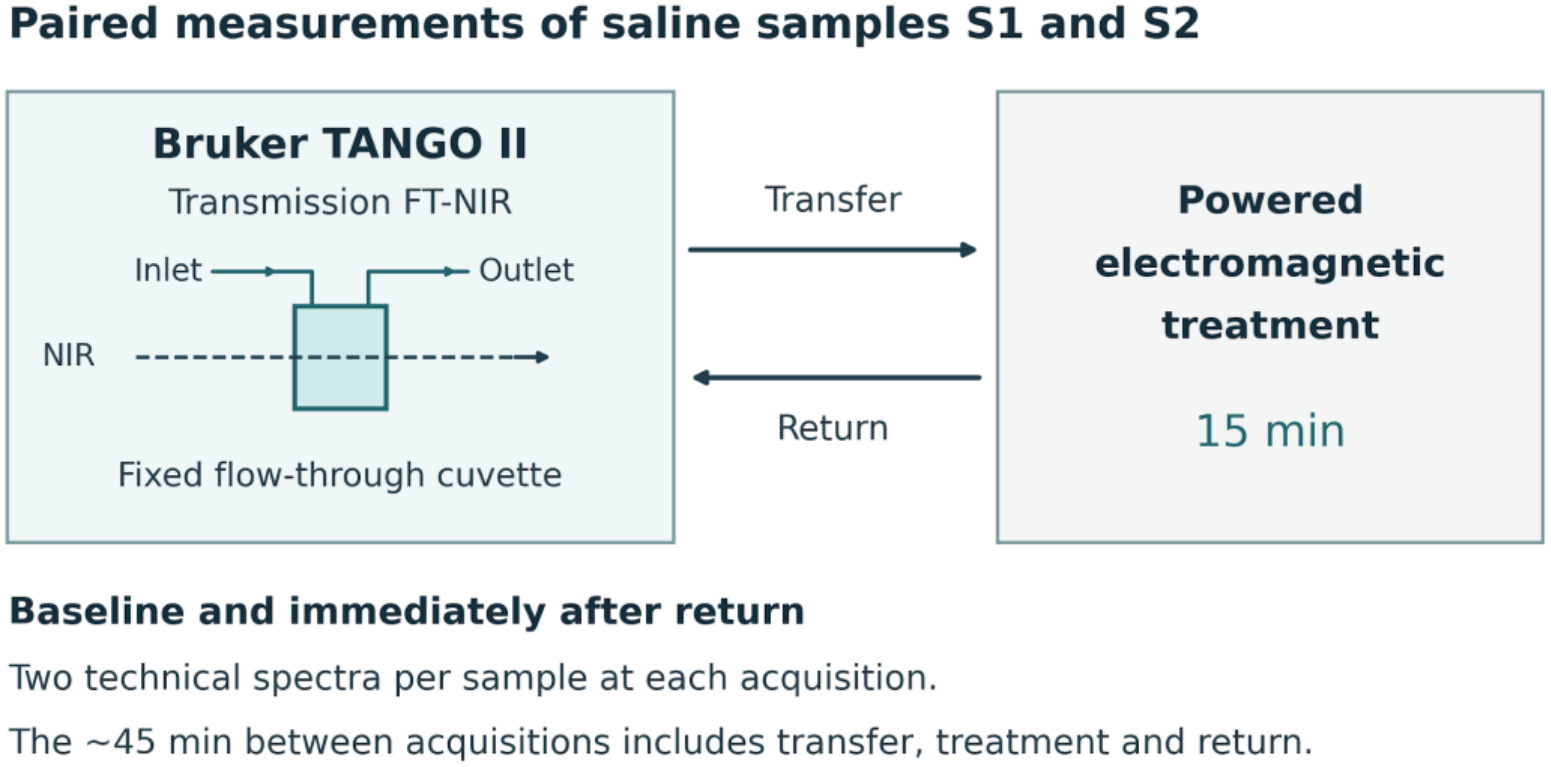
Experimental setup for the paired saline measurements. Each spectrum comprised 16 coadded scans. After scatter correction, the two technical spectra at each acquisition were averaged to give one pre/post difference spectrum per sample.

Transmission spectra were acquired with a Bruker TANGO II using 16 coadded scans and 8 cm^−1^ nominal resolution. OPUS StrayCorr-processed spectra contained 1900 points at 4 cm^−1^ spacing, spanning 11,544–3948 cm^−1^. Analysis used the 361 native points within the 1300–1600 nm first O–H overtone window. “Raw absorbance” here denotes the instrument-processed signal before the additional correction below. Separate ultrapure-water measurements were excluded from paired treatment differences and the preprocessing reference.

Multiplicative scatter correction (MSC) used the grand mean of the eight saline spectra as the common reference. Each spectrum j was fitted by ordinary least squares over spectral points k, then corrected as

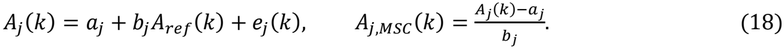

Corrected technical duplicates were averaged within each sample and condition before calculating post-treatment minus initial differences. The 12 water matrix coordinates (WAMACs) were sampled at native wavelengths nearest selected representative values within the conventional WAMAC ranges [13, 14]. All coordinates are reported, without interpolation or band integration. MSC differences describe relative spectral shape after fitted offset and scaling correction; they are not absolute concentrations.

### 4.2 Spectral redistribution and hydrogen bonding

Both pairs showed higher raw absorbance after the treatment sequence (Figure 3a). Within the analysis window, affine fits of post-treatment to initial mean spectra gave offsets of 0.05419 and 0.05012 absorbance units and slopes of 1.03085 and 1.02572 for S1 and S2, respectively. Examination after MSC resolves the smaller shape changes within this substantial offset and scaling response.

**Figure 3.**
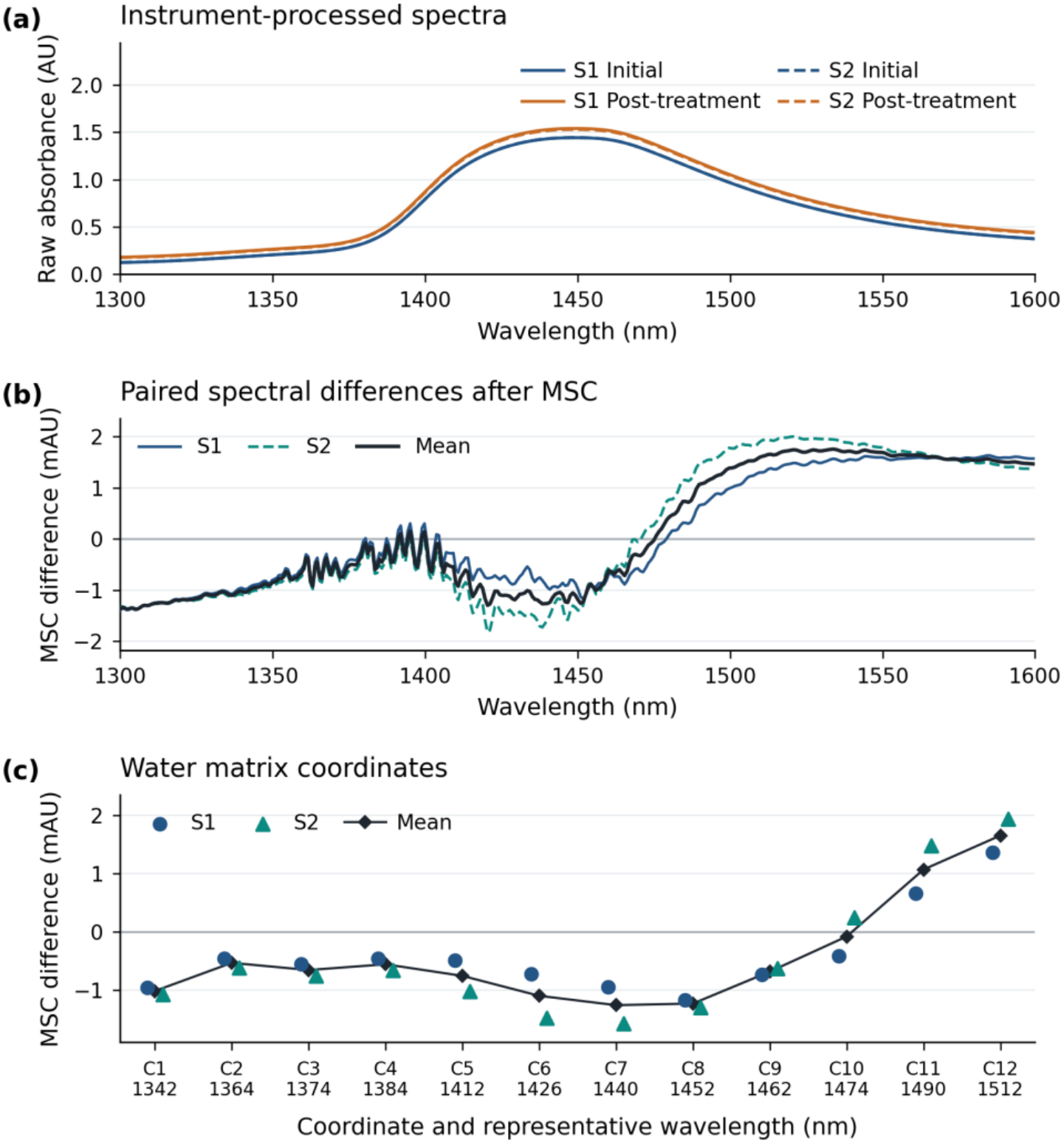
Preliminary FT-NIR response of seawater-derived saline solution. Two paired samples (S1, S2) were measured with two technical spectra per condition. (a) Mean instrument-processed absorbance before and after treatment. (b) Paired post-treatment minus initial differences after multiplicative scatter correction (MSC), with their mean. (c) Differences at the nearest native spectral points to 12 representative water matrix coordinates; wavelength labels identify the selected representative coordinates. Individual sample values are shown without inferential error bars. Differences are expressed in milli-absorbance units (1 mAU = 0.001 AU).

The paired MSC differences showed concordant signs at 11 of the 12 coordinates: C1–C9 decreased, C11–C12 increased, and C10 differed in direction between samples (Figure 3b–c and Table 1). Mean differences near 1440, 1452, 1490, and 1512 nm were −0.001262, −0.001234, +0.001065, and +0.001649 absorbance units. Showing both sample pairs makes the shared direction and sample variation visible without treating technical scans as additional biological or preparation replicates.

**Table 1.** All 12 WAMAC differences after MSC. Values are post-treatment minus initial in mAU; the mean is the arithmetic mean of S1 and S2. Wavelengths are the nearest native samples to the selected representative coordinates, without band integration.

| **WAMAC** | **Sampled wavelength (nm)** | **S1 (mAU)** | **S2 (mAU)** | **Mean (mAU)** |
|---|---|---|---|---|
| C1 | 1341.92 | -0.964 | -1.073 | -1.019 |
| C2 | 1363.88 | -0.456 | -0.615 | -0.536 |
| C3 | 1373.63 | -0.555 | -0.762 | -0.659 |
| C4 | 1384.27 | -0.457 | -0.663 | -0.560 |
| C5 | 1411.63 | -0.493 | -1.024 | -0.758 |
| C6 | 1426.13 | -0.724 | -1.477 | -1.101 |
| C7 | 1440.09 | -0.948 | -1.575 | -1.262 |
| C8 | 1451.80 | -1.172 | -1.296 | -1.234 |
| C9 | 1461.99 | -0.738 | -0.635 | -0.686 |
| C10 | 1474.06 | -0.412 | +0.245 | -0.084 |
| C11 | 1489.87 | +0.656 | +1.475 | +1.065 |
| C12 | 1512.40 | +1.361 | +1.937 | +1.649 |

In conventional aquaphotomic assignments, C7 and C9 include contributions associated with less extensively hydrogen-bonded environments, while C11 and C12 include four-hydrogen-bond and strongly bound water contributions [13, 14]. The relative increases at C11/C12, together with decreases at C5, C7, and C9, are therefore consistent with a shift toward a more extensively hydrogen-bonded or strongly bound water matrix. The assignments are tentative and overlapping; this interpretation concerns relative spectral contributions rather than counts of molecules or measured individual bond strengths.

The exploratory dataset contains two paired samples, with a mean temperature increase from 29.84 to 30.14 °C and initial acquisitions preceding post-treatment acquisitions. Matched sham measurements are absent, so temperature, handling, fluid exchange, and measurement sequence remain alternative contributors. MSC does not isolate these effects. The finding supplies a positive empirical target for the proposed structural response, while its attribution to electromagnetic dressing requires controlled comparison. Acquisition after return to the spectrometer also means that the observed spectrum samples the post-treatment material state, rather than the in-resonator oscillation calculated in Section 3.

## 5 Biological implications

We conjecture that water’s central role in biological organization arises in part from its ability to couple molecular polarization to the quantum electromagnetic vacuum as shaped by biological boundaries. In this view, cellular geometry, dielectric response, and membrane polarization modify the electromagnetic environment sampled by water, while collective water polarization contributes to that environment in return. If this interaction changes interfacial water correlations and hydration energetics, it could bias protein conformations and channel function. The proposed role of vacuum dressing therefore concerns a possible physical contribution to the established participation of water in biomolecular organization [1, 2, 22].

### 5.1 Hydration and functional plasticity

Protein function depends on the relative populations of conformational states and the rates of transitions between them. Hydration water contributes to these energetics, and molecular simulations show coupled fluctuations of protein side chains and surrounding water [22]. Equation (17) supplies a local mechanism by which a change in fast vibrational motion or fluctuations can alter the mean force on a slower structural coordinate. Extending this description to coupled hydration and protein coordinates would permit calculation of whether electromagnetic dressing shifts conformational free energies or transition barriers. Here, functional plasticity means a reversible redistribution among accessible functional states, rather than a general increase in rigidity or hydrogen-bond strength.

Protein channels provide a concrete test case. Molecular simulations of nearly 200 ion-channel structures show that pore radius and hydrophobicity influence water occupancy; local dewetting can create a barrier to ion permeation even without steric closure [23]. This establishes hydration as a functional variable in channel gating. We propose that a sufficiently large change in interfacial water response could, in turn, alter pore wetting, the stabilization of charged groups, or the free-energy balance between conducting and nonconducting states. Such effects would connect vacuum-modified hydration to channel activity, provided that their magnitude competes with thermal fluctuations and other gating forces. Figure 4 summarizes this proposed connection between the electromagnetic environment, hydration response, and conformational energetics.

**Figure 4.**
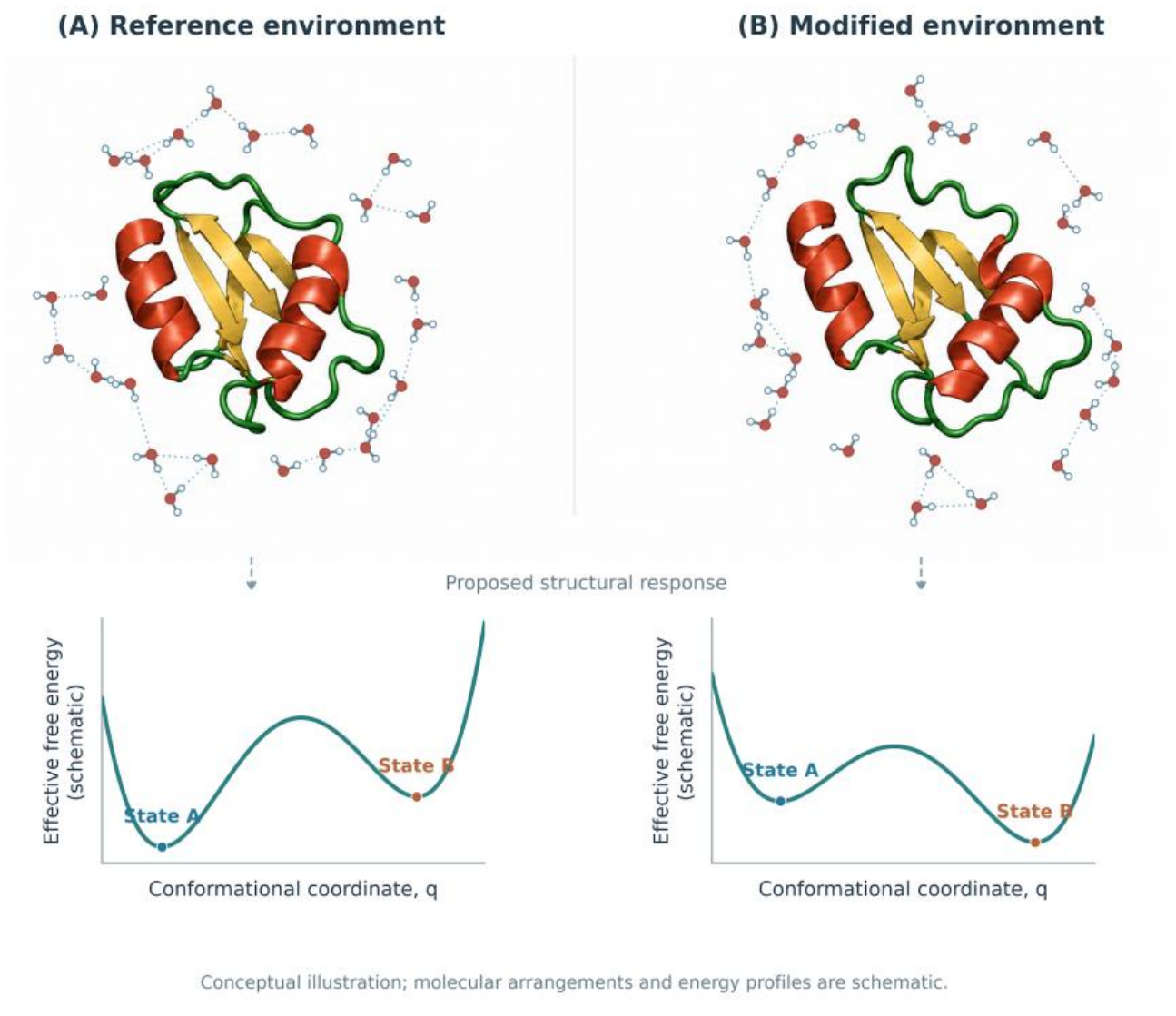
Proposed hydration-mediated conformational plasticity. Hydrated proteins and hydrogen-bond networks are shown schematically in a reference environment (A) and a modified environment (B). Dashed arrows indicate the hypothesized influence of collective water response on protein dynamics. The lower panels show illustrative effective free-energy profiles along a conformational coordinate *q*; changes in basin depths and barrier heights represent possible influences on state populations and transition kinetics. Molecular arrangements and energy profiles are schematic. The direction and magnitude of the depicted changes have not been calculated or established experimentally.

### 5.2 Biological resonators and mitochondrial interfaces

Energized membranes provide structured biological environments in which this hypothesis can be examined. Individual mitochondrial cristae exhibit distinct membrane potentials [24], and proton-driven membrane models predict changes in crista shape [25]. An illustrative 0.15 V drop across 5 nm gives an average transmembrane field of 3 × 10^7^ V m^−1^, although the field in adjacent hydration layers requires a spatially resolved calculation. Membrane bias and geometry could jointly modify interfacial susceptibility and the reaction field experienced by water.

Recent mitochondrial spectroscopy offers an experimentally motivated candidate for endogenous light–matter coupling. In a preprint, Yang, Gu, and Song report a feature near 71 THz associated with intact mitochondria, together with increased cellular ATP content following mid-infrared illumination [26]. Their model couples lipid CH_2_ vibrations near 87 THz to an organelle-scale electromagnetic mode and predicts branches near 71 and 103 THz. Cristae supply ordered lipid oscillators within this model; the proposed optical dimension is associated with the whole mitochondrion, rather than the nanometre spacing between cristae. These observations motivate investigation of biological resonators, while the assignment to vibrational polaritons remains a proposed interpretation. The study does not directly measure vacuum dressing of water.

The extension proposed here is that, where a biological structure supports suitable electromagnetic modes, adjacent water could participate through its vibrational dipoles and collective polarization. This requires a quantitative account of mode profiles, molecular overlap, dispersion, and losses. A narrow aqueous compartment need not constitute a conventional standing-wave cavity to modify the local electromagnetic response; its effect can instead be evaluated through the projected Green tensor and reaction kernel. Metabolic activity and electrochemical gradients supply explicit nonequilibrium inputs, while the vacuum contribution is assessed through the coupled fluctuation spectrum described in Section 2.3.

### 5.3 Coupling across scales

In our forthcoming contribution to the Springer proceedings of the 2025 Aquaphotomics conference in Kobe [27], we propose a hierarchy of coupled oscillators in which a Planck-scale restoring interaction undergoes scale-dependent screening, motivating characteristic frequencies across molecular and biological assemblies. Building on this framework, we postulate that Planck-scale vacuum organization may be coupled across successive material scales to molecular hydration and biological organization, with water acting as a dynamically responsive intermediary. Within the present formulation, this conjecture acquires an operational meaning through environment-dependent field correlations, reaction kernels, and collective susceptibilities. Establishing the cross-scale connection requires a derivation linking the proposed fundamental organization to these effective response functions, together with coupling strengths and relaxation rates. The frequency-scaling hypothesis motivates that program; it does not by itself establish dynamical transfer between the scales.

### 5.4 Experimental implications

A focused test would combine fundamental infrared or dielectric characterization with FT-NIR measurements and an independent protein or channel assay. Geometry or membrane bias could be varied while controlling temperature, pH, ionic composition, and sample handling. The proposed mechanism predicts a relationship between changes in the electromagnetic response, changes in hydration-sensitive observables, and changes in conformational populations or channel activity. Detuning and molecular-overlap dependence would help identify a resonant contribution; matched controls would be needed to separate it from direct field effects and heating. A vacuum-specific interpretation additionally requires the coupled fluctuation calculation described in Section 2.3. Correlation measurements would determine whether the effect extends beyond local hydration to collective biological organization.

## 6 Conclusion

This work links an environment-dependent electromagnetic response framework, a driven-resonator calculation, and preliminary FT-NIR observations of saline water. The model demonstrates tunable collective O–H amplitude and phase below resolved strong coupling. The paired spectra show redistribution consistent with greater contributions from extensively hydrogen-bonded or strongly bound water environments, providing an empirical target for controlled tests of hydration-mediated function.

The biological conjecture is that changes in water–vacuum coupling influence protein conformations and channel activity, potentially connecting fundamental vacuum organization to biological order. Establishing this connection requires a quantitative sequence from field–water coupling through structural free-energy changes to reproducible functional effects. Such tests would determine whether vacuum dressing contributes measurably to biological organization and functional plasticity.

## Notes

### Competing Interest Statement

The authors have declared no competing interest.

